# Competition between glycine and GABA_A_ receptors for gephyrin controls their equilibrium populations at inhibitory synapses

**DOI:** 10.1101/2024.12.09.627587

**Authors:** Dorota Kostrz, Stephanie A Maynard, Clemens Schulte, François Laurent, Jean-Baptiste Masson, Hans M Maric, Charlie Gosse, Antoine Triller, Terence R Strick, Christian G Specht

## Abstract

Glycine and GABA receptors are ligand-gated chloride channels that mediate inhibitory neurotransmission throughout the central nervous system. The receptors co-localise widely at inhibitory synapses in the spinal cord and in the brainstem due to their interaction with an overlapping binding site of the synaptic scaffold protein gephyrin, pointing to a direct competition between the different receptor types. We have put this hypothesis to the test using single molecule approaches to measure receptor-gephyrin interactions in cells and *in vitro*. We explored the effects of receptor competition at inhibitory synapses in living neurons by measuring the change in the accumulation and effective stabilisation energy of glycine receptors in the presence of interfering GABA receptor complexes through single molecule tracking and diffusion analysis. Secondly, using molecular tweezers, we quantified the thermodynamic properties of receptor-gephyrin binding, demonstrating direct and reversible competition through the addition of interacting peptides in solution. The relatively low affinity of GABA receptor subunits for gephyrin compared to the glycine receptor raises interesting questions about the role of this competition in synaptic plasticity. We hypothesize that GABA and glycine receptor competition constitutes a molecular system designed to reconcile synapse stability and plasticity at mixed inhibitory synapses.

## Introduction

Glycine receptors (GlyRs) and γ-aminobutyric acid type A receptors (GABA_A_Rs) are ligand-gated ion channels that mediate the majority of fast inhibitory synaptic transmission in the central nervous system. At many inhibitory synapses, the two receptor types act in concert. The neurotransmitters glycine and GABA are present in the same presynaptic vesicles ^1^, they are taken up by the common vesicular inhibitory amino acid transporter (VIAAT) and are co-released in the brainstem ^2, 3^, cerebellum ^4^ and in the spinal cord ^5^. The glycine uptake transporter GlyT2 and the GABA synthesising enzyme GAD are both present in certain inhibitory neurons in the brainstem and cerebellum that co-release GABA and glycine ^6^. In the spinal cord, postsynaptic GlyRs and GABA_A_Rs co-localise at mixed inhibitory synapses ^7^, where they mediate sensory and motor processing between the brain and periphery ^8^. Mixed GABAergic and glycinergic synapses represent 33-64% of inhibitory synapses in the spinal cord, depending on the lamina ^9, 10^. Mixed inhibitory transmission thus appears to play an important functional role at these synapses.

Neurotransmitter receptors diffuse laterally in the plasma membrane between extrasynaptic and synaptic sites, where they are transiently trapped either through binding to scaffold molecules or as a consequence of molecular crowding ^11^. In this way, the receptors are enriched in the post-synaptic membrane where their number determines synaptic strength. GlyRs and GABA_A_Rs share a common scaffolding protein, gephyrin. At the molecular level, gephyrin can bind the GlyR β-subunit as well as several of the GABA_A_R subunits at a common binding site ^12, 13^. Isothermal titration calorimetry (ITC) experiments have determined a dissociation equilibrium constant *K_D_*of 0.02 – 6 μM for the interaction of gephyrin with the intracellular loop of the GlyR β-subunit, and a *K_D_*of 17 μM and 5.3 μM for the GABA_A_R α1 and α3-subunits, respectively (^12, 13, 14, 15, 16, 17^, reviewed in ^18^). This raises the distinct possibility that the affinity of the receptors for gephyrin controls their mobility and recruitment at synapses. Direct competition between GlyRs and GABA_A_Rs for gephyrin-binding sites at synapses has not been formally demonstrated, but it is likely to be critical for determining the relative abundance and consequently the contribution of glycinergic and GABAergic transmission to inhibitory synaptic strength.

Competition between GlyR and GABA_A_R complexes is also likely to play a role in inhibitory synaptic plasticity, since changes in gephyrin-binding may cause a shift in the dynamic equilibrium of the two receptors ^15, 16^. This mechanism could provide important insights into the regulation of receptor numbers and dynamics in an activity-dependent manner or in response to deficits of inhibitory neurotransmission in a pathological setting. For example, super-resolution correlative light and electron microscopy (SR-CLEM) of endogenous GlyRs in spinal cord tissue uncovered a stereotypical modular organisation of the receptors that was maintained in a hypomorphic mouse model of the neuromotor disease hyperekplexia ^19^, which could limit functional compensation through GABA_A_Rs ^20^. Also, GlyRs and GABA_A_Rs were shown to form partially overlapping nanodomains at mixed spinal cord synapses ^21^, pointing to a non-homogenous spatial distribution. However, it is not known how the subsynaptic domains (SSDs) of GlyRs and GABA_A_Rs are formed and maintained in view of receptor competition for the same synaptic binding sites.

In this study, we combined two different single-molecule techniques to examine how GlyRs and GABA_A_Rs compete for gephyrin binding sites at mixed inhibitory synapses in cultured neurons and *in vitro*. Single molecule tracking of receptor diffusion showed a reduction in the effective stabilisation energy of GlyRs in the presence of GABA_A_Rs at spinal cord synapses, demonstrating competition between GlyRs and GABA_A_Rs in living neurons. Magnetic tweezer experiments further showed that GABA_A_R peptides corresponding to the gephyrin binding region of the α3-subunit effectively compete with the GlyR β-subunit for a common binding site of gephyrin, despite their lower thermodynamic stability.

## Results

### GABA_A_R α3 variants compete with GlyRs at mixed inhibitory synapses

To study the interplay between GlyRs and GABA_A_Rs at inhibitory synapses we chose cultured spinal cord neurons as a suitable cellular model that can be examined using single-molecule imaging approaches. The majority of inhibitory synapses in these neurons are mixed glycinergic and GABAergic synapses ^10^. Among the different GABA_A_R subunits we focussed on GABA_A_R α3, since it is widely expressed throughout the spinal cord ^22^ and because its intracellular loop displays the highest affinity for gephyrin *in vitro* ^13, 17^. We therefore anticipated an efficient competition of GABA_A_R α3 with GlyR β-subunits for synaptic binding sites in spinal cord neurons.

To explore how gephyrin-binding modulates GABA_A_R localisation at synapses, we selected two previously described variants of the wild-type GABA_A_R α3-subunit (GABA_A_R α3^WT^) that alter the receptors’ affinity to gephyrin: the high affinity variant GABA_A_R α3^N369S^, and the binding deficient variant GABA_A_R α3^F368A/I370A^ ^13^ (**Fig. 1a**). We hypothesised that increasing or decreasing the affinity of the GABA_A_R α3 constructs for gephyrin should induce bi-directional changes in the synaptic localisation of the receptor complexes. The overexpression of the recombinant GABA_A_R α3-subunits in spinal cord neurons is limited due to the fact that their surface expression requires co-assembly with endogenous GABA_A_R subunits ^23^.

**Figure 1.**
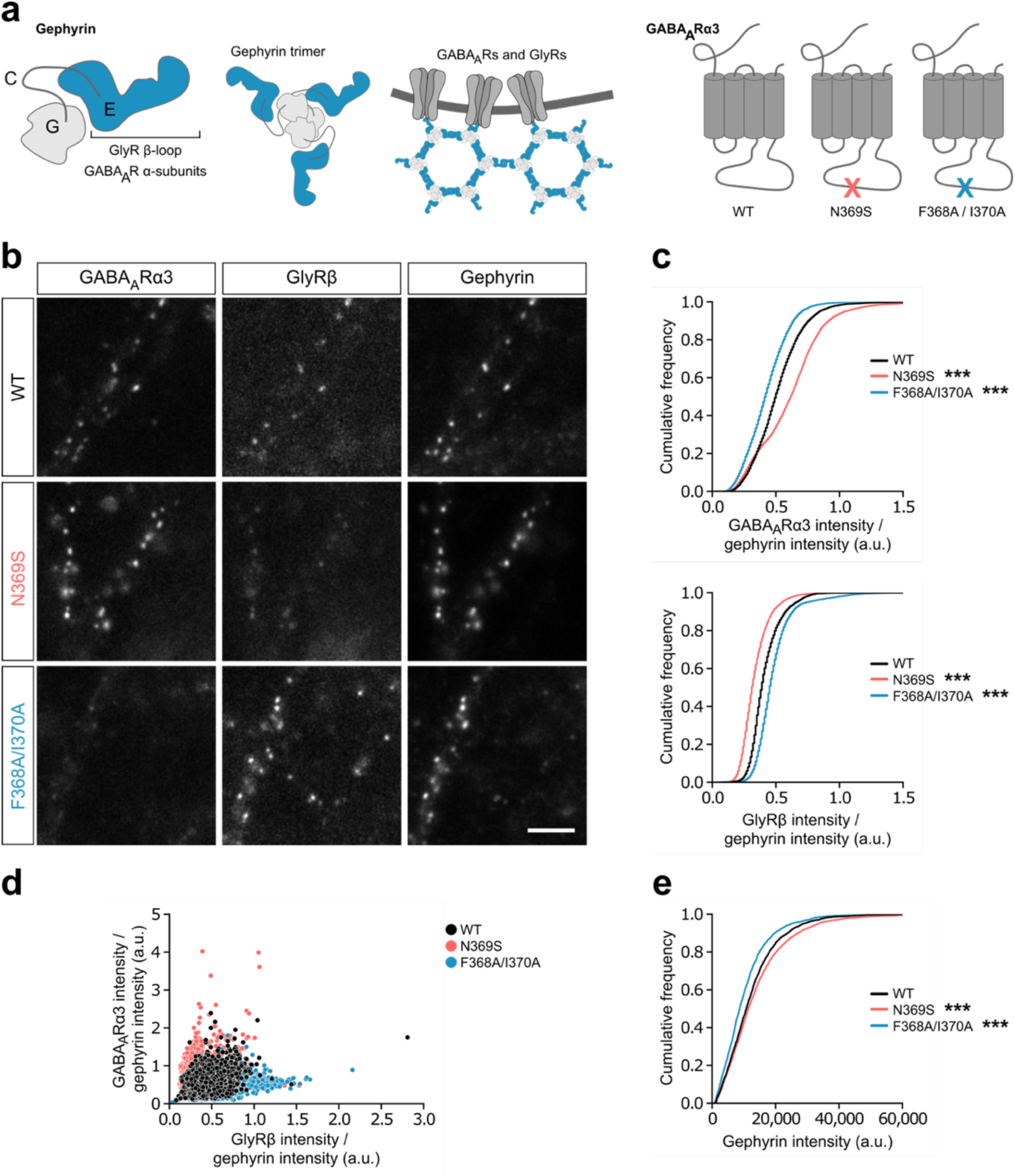
GlyR and GABA_A_R enrichment at synaptic gephyrin clusters at inhibitory synapses. (**a**) From left to right: Schematic of gephyrin domains, gephyrin trimer structure, receptor-gephyrin clustering at postsynaptic domain, and GABA_A_R-ɑ3 subunits with positions of gephyrin-binding mutants. (**b**) Representative images of GlyRs and α3-subunit containing GABA_A_Rs at inhibitory synapses. Lentiviruses were used for the expression of mScarlet-tagged GABA_A_R α3^WT^, α3^N369S^ (increased affinity), and α3^F368A/I370A^ (binding-deficient) in spinal cord neurons from mEos4b-GlyR β knock-in animals, followed by immunolabelling of gephyrin (mAb7a antibody). Scale bar = 5 µm. (**c**) Cumulative frequency distribution of the relative intensities of GABA_A_R α3 and GlyR β at gephyrin puncta in neurons expressing mScarlet-tagged GABA_A_R α3^WT^, α3^N369S^, or α3^F368A/I370A^, as well as mEos4b-GlyR β. GlyR and GABA_A_R levels were normalised to the gephyrin immunofluorescence intensity. (**d**) Scatter plot comparing the normalised GABA_A_R α3 and GlyR β intensities at individual gephyrin-mAb7a positive puncta. (**e**) Cumulative frequency distribution of gephyrin intensities. N = 30 images per condition from 3 independent experiments. Non-parametric Kruskal-Wallis with Dunn’s multiple comparison test. ***p<0.001.

We first assessed the ability of the mutant receptors to accumulate at mixed inhibitory synapses and to alter GlyR occupancy at the gephyrin scaffold. Lentiviral constructs driving the expression of fluorescent mScarlet-tagged wild-type and mutant GABA_A_R α3-subunits were used to infect spinal cord neuron cultures expressing endogenous mEos4b-tagged GlyR β-subunits (allele *Glrb^tm^*^1^*^(Eos^*^4^*^)Ics^*, accession number MGI:6331065, ^19^). The neurons were then fixed and immunolabelled with mAb7a antibody against endogenous gephyrin to visualise inhibitory synapses using conventional fluorescence microscopy (**Fig. 1b**). We measured the intensity of mEos4b-GlyR β and mScarlet-GABA_A_R α3 fluorescence relative to gephyrin at individual synapses. In these imaging conditions, the mEos4b photoconvertible protein is not exposed to UV illumination, remaining in its green fluorescent form instead of being converted into the red form (**Supplementary Fig. S1**).

As expected, the normalized GABA_A_R α3^N369S^ fluorescence was increased at synapses compared to the GABA_A_R α3^WT^-subunit. The expression of GABA_A_R α3^N369S^ also substantially decreased the mEos4b-GlyR β intensity, suggesting that the accumulation of GlyRs is affected by the presence of the high affinity GABA_A_R variant due to direct competition (**Fig. 1c**). Conversely, expression of the binding-deficient GABA_A_R α3^F368A/I370A^ led to a reduction in normalized fluorescence compared to GABA_A_R α3^WT^ and an increase in mEos4b-GlyR β intensity (**Fig. 1c** and **d**). This clustering behaviour is in agreement with the expected competition of the receptors for gephyrin-binding sites at mixed inhibitory synapses. Interestingly, gephyrin itself was also increased in the presence of GABA_A_R α3^N369S^ and decreased in the presence of GABA_A_R α3^F368A/I370A^ (**Fig. 1e**), an observation consistent with the reciprocal stabilisation of gephyrin through receptor binding. The data thus confirm that α3-containing GABA_A_Rs accumulate at mixed inhibitory synapses in accordance with the strength of their gephyrin-binding domains, pointing to a direct competition with endogenous GlyRs.

### High affinity GABA_A_R α3 variants alter the GlyR trapping potential at synapses

The previous findings show that GABA_A_Rs containing the recombinant α3^N369S^-subunit affect GlyR clustering at mixed inhibitory synapses. To provide a spatiotemporal description of the diffusion dynamics of the receptor complexes in cultured spinal cord neurons, we made use of single particle tracking (SPT) based on photoactivated localisation microscopy (PALM) of endogenous GlyRs in the synaptic and extrasynaptic plasma membrane. sptPALM generates high-density datasets of hundreds of thousands of protein localisations over short periods of time, which lends itself to statistical analysis to determine the effective diffusivity and trapping potential of biomolecules in living cells ^24, 25^.

GlyR mobility was studied in living neurons expressing endogenous mEos4b-GlyR β that had been infected with lentivirus constructs of Halo-tagged GABA_A_R α3-subunits (constructs Halo-Gabra3^WT^ and Halo-Gabra3^N369S^, see **Methods**). The expression of the GABA_A_R constructs was verified through the addition of the fluorescent JF646-Halo-ligand (**Supplementary Fig. S2**). In line with the epifluorescence data (**Fig. 1**), the number of mEos4b-GlyR β detections per synapse was lower in the presence of the GABA_A_R α3^N369S^ variant compared to α3^WT^ (**Fig. 2a** and **b, Supplementary Fig. S3**), again showing the loss of GlyRs at synapses due to competition. SPT trajectories were recorded by following the movement of mEos4b-GlyR β containing GlyRs with a temporal resolution of 15 ms (67 Hz streamed acquisition, **Fig. 2c**).

**Figure 2.**
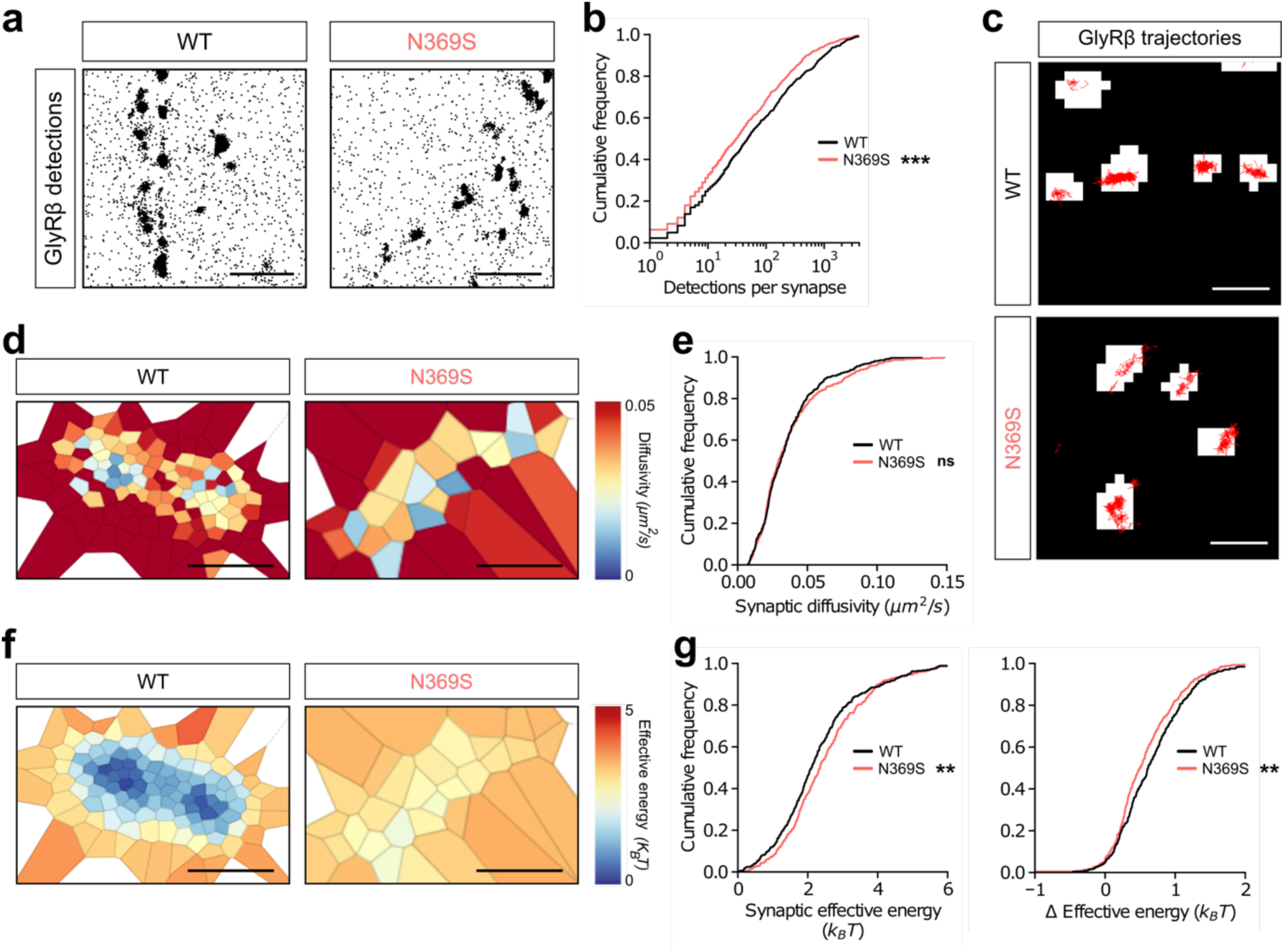
Effects of GABA_A_Rs on diffusion dynamics and effective potential trapping energy of GlyRs at synapses. (**a**) Representative pointillist images of mEos4b-GlyR β detections at mixed inhibitory spinal cord synapses expressing Halo-tagged GABA_A_R α3^WT^ or α3^N369S^ labelled with JF646-Halo-ligand. SPT movies of mEos4b-GlyR β of 40,000 frames were recorded in living neurons at 67 Hz. Scale = 2 µm. (**b**) Cumulative frequency distribution of the number of mEos4b-GlyR β detections per synapse shows fewer GlyRs at synapses expressing Halo-GABA_A_R α3^N369S^. (**c**) GlyR trajectories at individual synapses expressing Halo-GABA_A_R α3^WT^ or Halo-GABA_A_R α3^N369S^. Scale bar = 500 nm. (**d**) Examples of Voronoi tessellation showing the diffusivity at an individual synapse expressing Halo-tagged GABA_A_R α3^WT^ or α3^N369S^. Scale bar = 100 nm. (**e**) The cumulative frequency distribution of the diffusivity of mEos4b-GlyR β at synapses expressing Halo-GABA_A_R α3^WT^ or α3^N369S^ shows that diffusion properties of GlyR are not altered by changes in affinity and/or the effective concentration of GABA_A_Rs. (**f**) Voronoi tessellation plots of the effective trapping energy at an individual synapse expressing Halo-GABA_A_R α3^WT^ or α3^N369S^. Scale bar = 100 nm. (**g**) Cumulative frequency distribution of the effective energy at synapses (left) and the change Δ in effective energy of mEos4b-GlyR β between the synaptic and extrasynaptic regions (right) show that the GlyR is less stabilised in the presence of the high-affinity Halo-GABA_A_R α3^N369S^ variant. N = 34 movies per condition from 3 independent experiments. Non-parametric unpaired two-tailed t-test, with Mann-Whitney post hoc. ***p<0.001, **p<0.01, ns = not significant.

The effect of α3^N369S^-containing GABA_A_Rs on GlyR dynamics was analysed using Bayesian inference, so as to derive physical parameters that control receptor motion at synapses ^26^. Based on changes observed in GlyR localisation density maps we found that the effective diffusivity of GlyRs at synapses was not affected by the presence of the GABA_A_R α3^N369S^ mutant (**Fig. 2d** and **e**). We further quantified the effective stabilisation energy of the GlyRs as a measure of receptor trapping at synapses (**Fig. 2f** and **g**). We observed that the expression of the Halo-GABA_A_R α3^N369S^ variant increased the effective energy of GlyRs at synapses, thus decreasing the depth of the potential well that is the difference (Δ) between the effective energy inside the synaptic domain compared to the extrasynaptic plasma membrane (**Fig. 2g**). This result points to a direct competition between the two types of receptor, where a higher occupancy of synaptic binding sites by GABA_A_R α3^N369S^ shifts the equilibrium towards a decreased effective dwell time of GlyRs at synapses.

### GlyR β-loop binds reversibly to gephyrin with a residence time of tens of seconds and an affinity in the nanomolar range

To quantitatively describe the phenomena observed in living cells, we developed a nanomechanical *in vitro* assay in which a gephyrin trimer and the cytosolic β-loop of GlyR were fused to SNAP- and CLIP-tags, respectively (**Supplementary Fig. S4 and S5**), and covalently engrafted at the tips of so-called junctured-DNA (J-DNA) forceps ^27^ (**Fig. 3a**). In this way, pairs of interacting molecules can be manipulated with magnetic tweezers and observed in real-time. Monitoring of the magnetic bead position under negligible force (*F* = 40 fN) allows one to detect if the nucleic acid scaffold is in a closed and compact conformation or in an open and extended one, i.e. if the protein complex is formed or not ^28, 29^ (**Fig. 3a**, upper panels, **and Supplementary Table S1**). By detecting a large number of transitions for a single pair of interacting molecules, we verified that, as expected for a stochastic process, the distributions of the dwell times in both conformations, Δ*t*_*closed*_ and Δ*t*_*open*_, were well fit by monoexponentials, hence providing time constants *τ*_*closed*_ and *τ*_*open*_(**Fig. 3b** and **c**, black curves). Next, comparing the data from 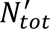 = 49 scaffolds, we found that the *τ*_*closed*_ and *τ*_*open*_values are themselves distributed normally around 22.7 ± 0.4 s and 68.5 ± 3.2 s (SE, **Supplementary Fig. S6a and b**), respectively. The dissociation rate constant of the GlyR-gephyrin complex being the inverse of the closed conformation characteristic time, we computed <*k_D_*> as 1/<*τ_closed_*> = 0.044 ± 0.005 s^-1^ (SEM, 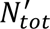 = 49 scaffolds; **Table 1**).

**Figure 3.**
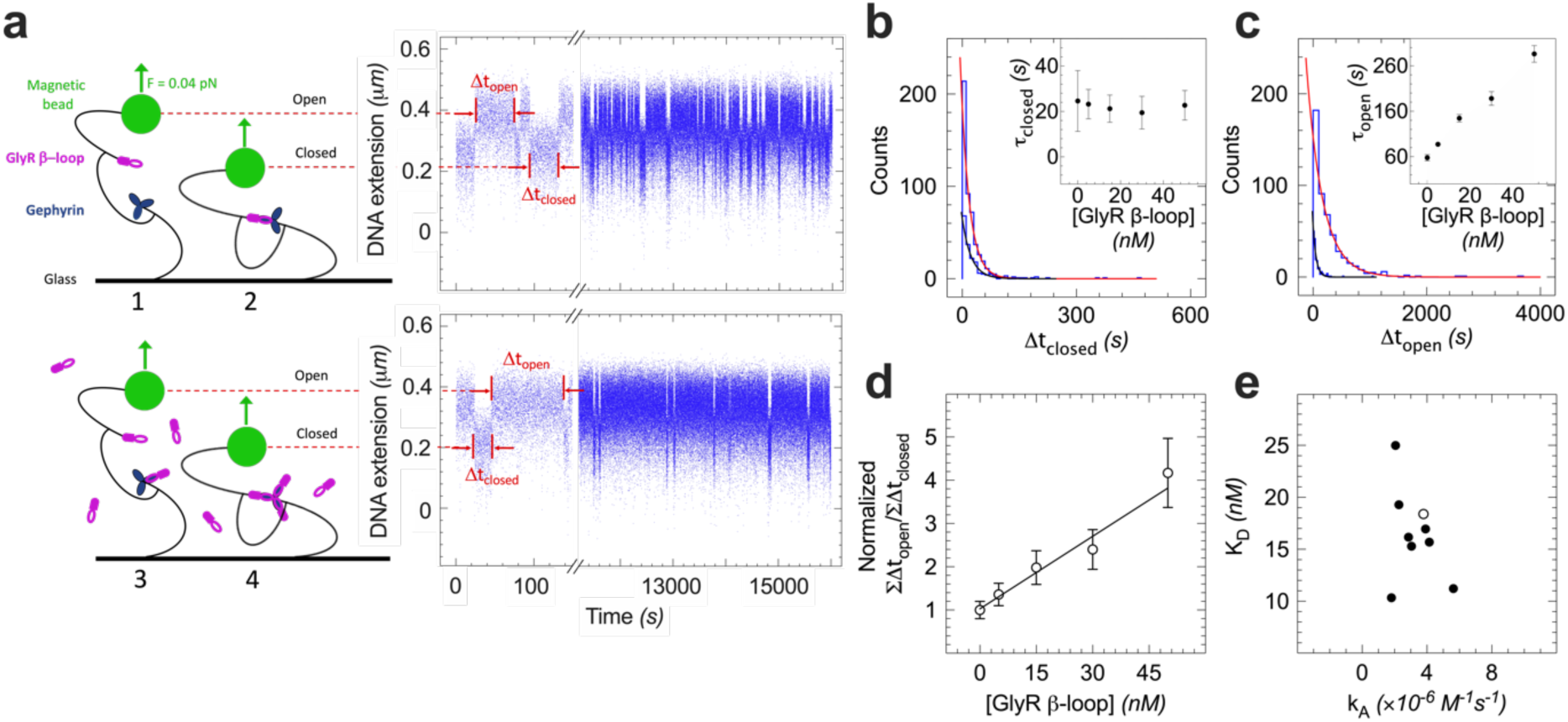
Titration by the GlyR β-loop of single J-DNA scaffolds engrafted with a gephyrin trimer and the GlyR β-loop. Measurements were conducted in gephyrin buffer at 19.2°C and under a constant force of 40 fN. (**a**) Representative time traces of the J-DNA extension showing spontaneous changes between closed and open conformations in the absence (top panel) and presence of 50 nM GlyR β-loop in solution (bottom panel). The period spent at low extension corresponds to the formation of an *in cis* complex between the tips of the forceps and is characterised by a dwell time Δ*t_closed_*. Conversely, in the absence of an interaction the extension is high (dwell time Δ*t_open_*). The addition of GlyR β-loop in solution enables various interactions *in trans* that do not modify the extension of the scaffold but influence its dynamics, as illustrated in the scheme (see also **Fig. S7**). (**b**) Single-scaffold Δ*t_closed_* histograms for [GlyR β-loop] = 0 and 50 nM. Fits to single-exponential distributions yield the respective characteristic times *τ_open_*= 24.6 ± 3.3 and 22.7 ± 1.5 s (SEM, *n* = 175 and 497 dwell times, black and red lines). Inset: Variation of *τ_closed_* as a function of [GlyR β-loop] for the shown J-DNA scaffold. (**c**) Same analysis for Δ*t_open_*, resulting in *τ*_open_ = 57.6 ± 7.0 and 286.3 ± 18.3 s for [GlyR β-loop] = 0 and 50 nM, respectively (SEM, *n* = 175 and 497, black and red lines. Inset: *τ_open_* as a function of the titrant. (**d**) Variation of the ratio between the total times that the scaffold spends in the open and closed conformation, Δ*t_open_* and Δ*t_closed_*, as a function of titrant concentration. Data were divided by the value measured in the absence of titrant and fit to a straight line, yielding *k_D_*= 18.5 ± 4.9 nM (SEM; see **Methods**). (**e**) Scatter plot of *k_D_* and *k_A_* constants determined for each of the 9 analysed J-DNAs, the open symbol referring to the scaffold shown in the previous panels (see **Text** and **Table 1** for the averaged values, **Fig. S8** for individual fits and **Table S2** for statistics). For clarity, the error values are not shown.

**Table 1.** Kinetic and thermodynamic parameters of the interaction between one of the three binding sites of the gephyrin trimer and the intracellular loops and peptides derived from GlyR β, GABA R α3^WT^, and GABA R α3^N369S^. Measurements were conducted in gephyrin buffer at 19.2°C. Values from single-molecule titration assays at constant force were obtained from the data shown in **Fig. 3c, 4e, S6c, and S8c** (see **Table S2** for statistics). Values from single-molecule dissociation assays with force step application come from **Fig. 5c** (see **Table S3** for statistics). Errors are given as SEM. na = not applicable, nd = not determined.

| Gephyrin partner | Length (aa) | Constant force experiments at 0.04 pN |  |  | Cycling force experiments at 1.1 pN |
| --- | --- | --- | --- | --- | --- |
| | | $\langle k_D \rangle$ (s<sup>-1</sup>) | $\langle K_D \rangle$ (nM) | $\langle k_A \rangle$ (10<sup>-6</sup> M<sup>-1</sup> s<sup>-1</sup>) | $k_D^{1.1 \text{ pN}}$ (s<sup>-1</sup>) |
| GlyR β-loop | 120 | $0.044 \pm 0.005^a$ | $16.5 \pm 4.0^b$ | $3.3 \pm 0.7^b$ | $0.059 \pm 0.003^c$ |
| GABA<sub>A</sub>R $\alpha 3^{\text{WT}}$-loop | 98 | nd | nd | nd | $71.4 \pm 5.1$ |
| GABA<sub>A</sub>R $\alpha 3^{\text{N369S}}$-loop | 98 | nd | nd | nd | $0.500 \pm 0.025$ |
| GlyR β-peptide | 11 | na | $2120 \pm 470$ | na | na |
| GABA<sub>A</sub>R $\alpha 3^{\text{WT}}$-peptide | 12 | na | $39400 \pm 17280$ | na | na |
| GABA<sub>A</sub>R $\alpha 3^{\text{N369S}}$-peptide | 12 | na | $7920 \pm 3420$ | na | na |

<sup>a</sup> Data collected over $N'_{\text{tot}} = 49$ scaffolds from all titration experiments in absence of titrant.
<sup>b</sup> Data collected over $N = 9$ scaffolds from the GlyR β-loop titration experiments at all GlyR β-loop concentrations.
<sup>c</sup> Averaged value from data collected over $N = 50$ scaffolds.

To obtain the bimolecular association rate constant, *k_A_*, and the dissociation equilibrium constant, *K_D_*, we relied on a titration protocol that introduces a well-controlled concentration parameter ^28, 30, 31^. Progressive addition of GlyR β-loop *in trans* (in solution) did not alter the duration of the closed conformation (**Fig. 3a,b**), which can be easily explained by the fact that, in absence of cooperativity, the dissociation *in cis* (the J-DNA engrafted gephyrin trimer and GlyR β-loop) is not influenced by the occupancy of the other two binding sites of the gephyrin trimer. In contrast, the time spent in the open conformation increased at higher concentrations of the GlyR β-loop, roughly quadrupling at 50 nM GlyR β-loop (**Fig. 3a,c**). This observation is in line with an association *in cis* that is impeded by the competing GlyR β-loop in the solution.

We further analysed the data by computing the ratio between the total amount of time that the scaffold spent in the open and in the closed conformation, ΣΔ*t_open_* ΣΔ*t_closed_*. As expected, this observable increases proportionally to the concentration of the titrant. After normalization by the value measured in the absence of GlyR β-loop in the buffer, the slope corresponds to 1*k_D_* (see **Methods and Supplementary Fig. S7**). Thus, for each gephyrin molecule a dissociation equilibrium constant of the GlyR β-loop could be determined (**Fig. 3d and Supplementary Fig. S8a**). Averaging over the entire scaffold population yielded *k_D_* = 16.5 ± 4.03 nM (SEM, *N* = 9; **Fig. 3e and Table 1**), close to the lowest values measured by ITC ^18^. Using the <*k_D_*> value determined above, we calculated the association rate constant for each individual J-DNA and obtained, after averaging over a population of N = 9, the mean association rate *k_A_* = (3.3 ± 0.7)·10^-6^ M^-1^s^-1^ (SEM, **Fig. 3e and Table 1**).

### GABA_A_R α3-peptides compete with GlyR β-loop for gephyrin binding sites

Having established the kinetic and thermodynamic properties of the high-affinity binding of the GlyR β-loop to gephyrin, we set out to interrogate the competition with the GABA_A_R α3-subunit. We used the titration assay described above, this time adding various interacting peptides in solution (**Fig. 4**). In this case, the functionalised scaffold works as a sensor reporting on the interactions taking place *in trans* (see **Methods**) ^28^.

**Figure 4.**
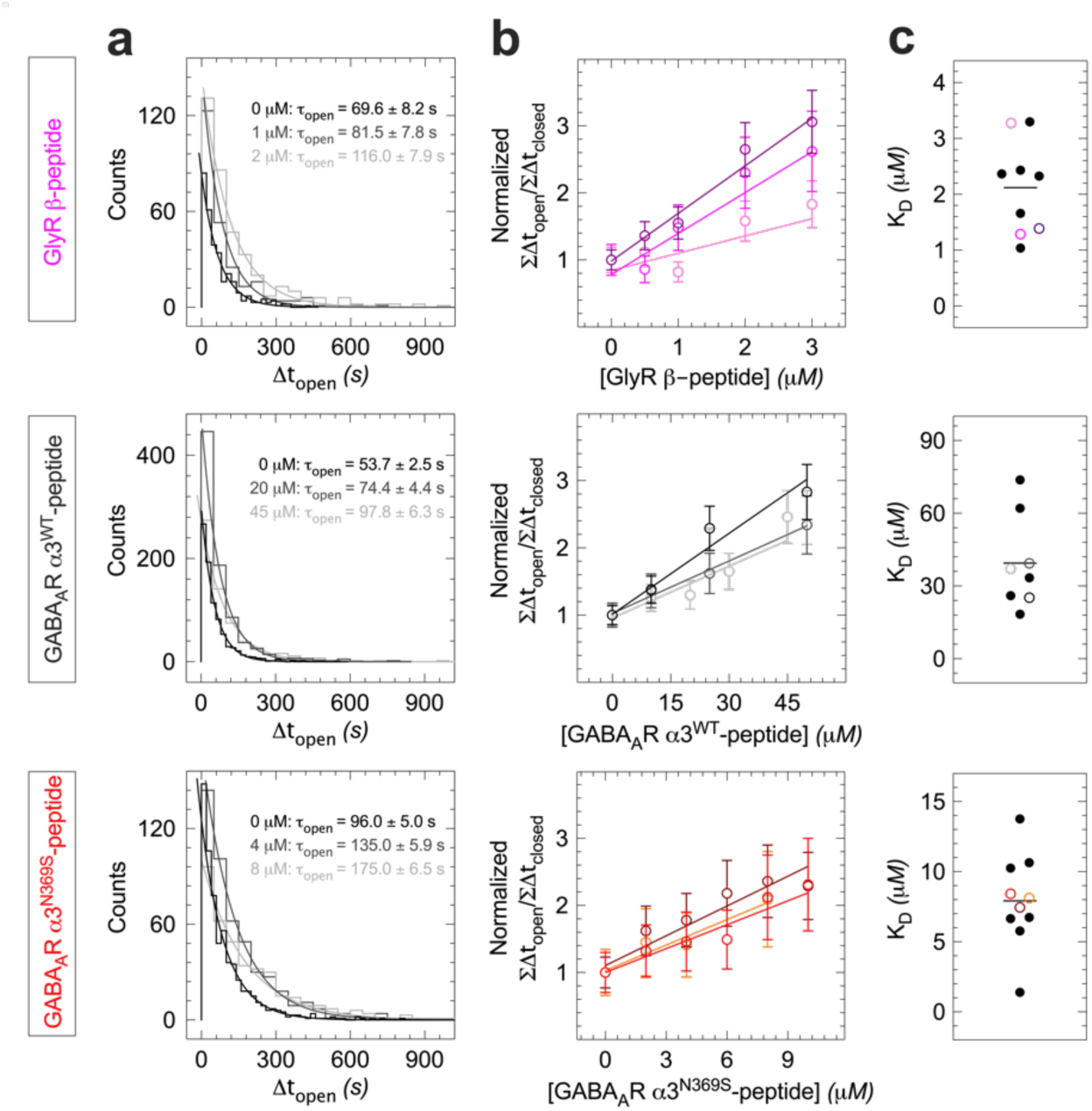
Titration of single J-DNA scaffolds engrafted with a gephyrin trimer and GlyR β-loop with peptides derived from GlyR β, GABA_A_R α3^WT^, and GABA_A_R α3^N369S^. Measurements were performed in gephyrin buffer at 19.2°C under a constant force of 40 fN and analysed as in Fig. 3. (**a**) Single-scaffold histograms for the dwell times in the open conformation, Δ*t_open_*. The acquisitions were performed at three different titrant concentrations: 0 (black), ≍ 0.5 *k_D_* (dark grey), and ≍ 1 *k_D_*(light grey), and the data fit to single-exponential distributions to obtain the characteristic time *τ_open_*. (**b**) Variation of the ratio between the total times of the open and closed conformations as a function of titrant concentration for three individual scaffolds per condition. (**c**) Dispersion of the *k_D_* values of all measured scaffolds; open symbols refer to the three scaffolds shown in **b** and horizontal lines mark the average values (see **Text** and **Table 1** for average values, **Fig. S9** for individual fits and **Table S2** for statistics). For clarity, the error values are not shown.

We first added soluble GlyR β-peptide corresponding to the core binding sequence of the intracellular loop of the GlyR β-subunit (amino acid residues F398 to F408). In agreement with our findings using the full-length β-loop as titrant, competition increased the values of *τ_open_* (**Fig. 4a**, top) and the normalized ΣΔ*t_open_*/ΣΔ*t_closed_* ratio (**Fig. 4b**, top). Linear fitting of the titration plots provided individual dissociation equilibrium constants for the binding of the GlyR β-peptide to gephyrin (**Fig. 4c**, top, **and Supplementary Fig. S9**), with *k_D_* = 2.12 ± 0.47 μM (SEM, *N* = 9; **Table 1**). The 100-fold higher value compared to the full-length GlyR β-loop is likely due to the absence of the regions flanking the core peptide, which are known to contribute to the interaction ^32^. Besides demonstrating that the competing molecules occupy the same binding pocket as the GlyR β-loop, these experiments provide quantitative data on the thermodynamic stability of the complex formed between the titrant and gephyrin.

Next, we added *in trans* the peptide corresponding to the gephyrin-binding sequence of the cytoplasmic loop of the GABA_A_R α3^WT^-subunit (**Fig. 4a-c**, middle panels). We determined a higher *k_D_* of 39.41 ± 17.28 μM (SEM, *N* = 8; **Supplementary Fig. S9 and Table 1**), in line with the comparably weak ability of wild-type GABA_A_Rs to displace GlyRs at synapses. For the α3^N369S^ variant of the same peptide we found a *k_D_* = 7.92 ± 3.42 μM (SEM, *N* = 10; **Fig. 4a-c**, bottom panels, **Supplementary S9, and Table 1**). This value is closer to the one obtained for the GlyR β-peptide, which reflects the ability of GABA_A_R complexes containing the α3-subunit with the N369S mutation to compete more strongly with GlyRs in cellular assays.

### The lifetimes of receptor-gephyrin complexes differ by two orders of magnitude

Although the titrations carried out with the core peptides suggest that the wild-type GABA_A_R α3-gephyrin complex has a higher dissociation equilibrium constant than the GlyR β-gephyrin complex, it is uncertain whether this is due to a lower association rate constant, a higher dissociation rate constant, or a combination of the two. To answer this question, we used a modified version of our nanomechanical single-molecule assay: gephyrin trimers and receptor loops were engrafted onto the J-DNA and the force was cycled between 0.001 pN, a low value at which the scaffolded partners are allowed to associate, and 1.1 pN, a higher value at which the Δ*t_closed_* dwell time before dissociation is measured (**Fig. 5a**). Interactions were detected as intermediate plateaux or shoulders between the low extension signal corresponding to both the open and the closed conformations at low force and the high extension signal corresponding to the open conformation at high force ^27, 29, 33, 34^ (**Fig. 5b**). Owing to the high temporal resolution of the camera, single binding events having a duration as short as 55 ms could be spotted (**Fig. 5b**, middle). Again, the Δ*t_closed_* values were exponentially distributed, and a fit to the histogram led to a characteristic time, 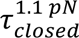 denoting the mean lifetime of the complex at the applied high force (**Fig. 5c**). It should be noted that in this protocol the change in extension observed upon dissociation, *Δz*, is fully determined by the geometry and mechanics of the J-DNA scaffold, as well as by the applied force. Events that did not comply with the expected change in extension could thus be attributed to non-specific interactions and excluded (**Fig. 5c**, insets, **and Supplementary Table S3**).

**Figure 5.**
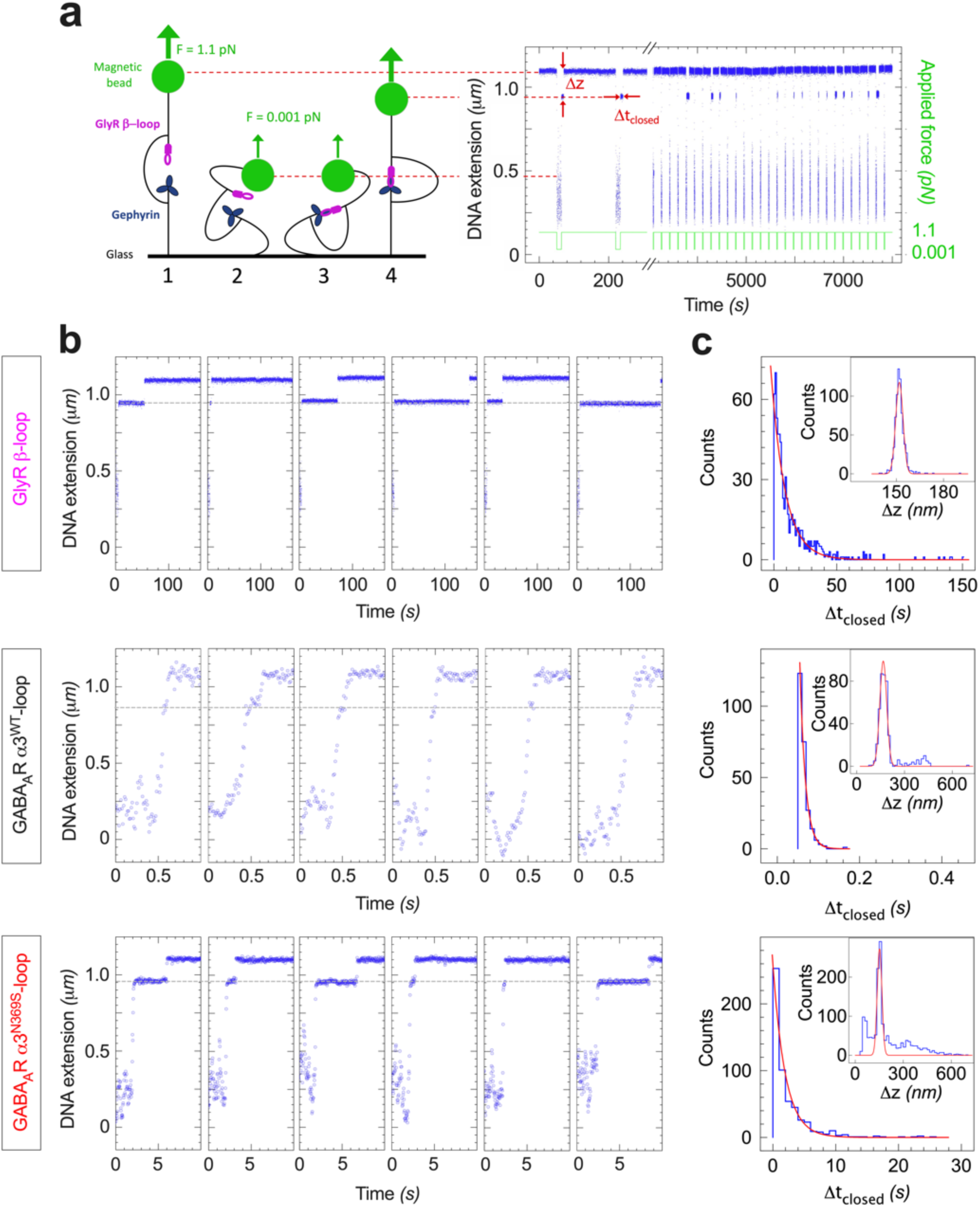
Lifetimes of the interaction between gephyrin and the intracellular loops of GlyR β, GABA_A_R α3^WT^, and GABA_A_R α3^N369S^ under an applied force of 1.1 pN. (**a**) Typical time trace of J-DNA forceps engrafted with a gephyrin trimer and the GlyR β-loop, the force alternating between 0.001 pN for 13 s and at 1.1 pN for 155 s. Starting at high-force with a dissociated complex (1) the force is reduced, which lowers the DNA extension and brings the tips into proximity (2), allowing the proteins to bind (3). The force is then switched back to 1.1 pN and the applied tension holds the scaffold in an intermediate extension which reports on the association state of the two partners (4). The J-DNA eventually recovers its initial high-extension as the complex dissociates (1). Each interaction is characterized by its duration, Δ*t_closed_*, and by the resulting change in extension observed upon dissociation, *Δz*. Blue points are raw extension data obtained at 31 Hz, while the green trace depicts the force modulation. (**b**) Individual interactions observed for the GlyR β, GABA_A_R α3^WT^, and GABA_A_R α3^N369S^-loop constructs (from top to bottom). Dashed lines indicate the expected high-force extension when a complex is formed. (**c**) Histograms of the Δ*t_closed_* dwell times. Top panel shows data from a single J-DNA scaffold with GlyR β-loop, fitted to single-exponential distribution and yielding 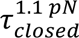=11.3 ± 0.6 s, while the mean value over N = 50 scaffolds was 16.9 ± 0.8 s (see **Fig. S13**). Data from different J-DNA scaffolds with GABA_A_R α3^WT^ and GABA_A_R α3^N369S^-loops were pooled before fitting to single-exponential distributions and yielded 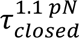 of 0.014 ± 0.001 s and 2.0 ± 0.1 s, respectively (see **Table 1** for the corresponding dissociation rate constants at 1.1 pN, 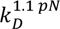, and **Table S3** *z* extension changes and fit to Gaussian distributions, yielding 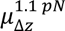 ranging from 152.0 to 166.6 nm (see **Methods** and **Table S3** for results and statistics). Measurements were conducted at 19.2°C in gephyrin buffer.

For the GlyR β-loop interaction with gephyrin at 1.1 pN, we determined 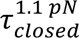 = 16.9 ± 0.8 s by averaging the characteristic times collected over *N* = 50 individual J-DNA scaffolds (SEM, see **Fig. 5c**, top, for a single scaffold, and **Supplementary Fig. S13** for the 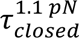 distribution). This value is only about a factor of two shorter than the one measured at a constant force of 0.04 pN (**Fig. 3b and Supplementary Fig. S6a**). This points to a weak force-dependence of the interaction, as confirmed by measurements at up to ≈ 10 pN (**Supplementary Fig. S10**). A fit to the Bell equation provided a 7.1 ± 0.4 Å Bell parameter and a dissociation characteristic time at zero-force 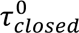 = 20.1 ± 1.1 s (SEM). It also shows that the dissociation phenomena observed in the low fN range can be considered identical to what would be seen in solution. Next, we probed the binding of full-length GABA_A_R α3^WT^-loop to gephyrin. We observed much shorter-lived complexes with 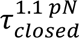 = 0.014 ± 0.001 s (SEM, *n* = 251, *N* = 22; **Fig. 5c** middle). Finally, the GABA_A_R α3^N369S^-loop interacted with gephyrin with a lifetime equal to 2.0 ± 0.1 s (SEM, *n* = 552, *N* = 20; **Fig. 5c**, bottom). Thus, the 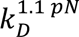 = 1/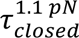 rate constants of the cycling-force assay show a similar pattern as the average equilibrium constants *k_D_* obtained from the titrations at constant force (**Table 1 and Supplementary Fig. S11**). We conclude that the differences in affinity between the different variants are essentially accounted for by differences in their interaction lifetimes.

Upon examination of the entire dataset, three unusual phenomena were observed, either at the single time-trace or at the J-DNA population levels. First, even though a majority of individual GlyR β-loop-gephyrin complexes displayed a constant extension change throughout the acquisition cycles, about 20% exhibited a bimodal distribution of the extension jump, Δ*z* (**Supplementary Fig. S12**). For these 12 scaffolds (from a population of N = 50), between 3.1% and 88.3% of the dissociations had additional, shorter extension jumps. The separation between the peaks ranged from 2.6 to 29 nm (**Supplementary Fig. S12c-e**). The events of both peaks were exponentially distributed and their characteristic times 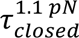 did not differ significantly (**Supplementary Fig. S12b**). The difference in Δ*z* could be the result of conformational changes within gephyrin ^35^ or spatial rearrangements of the gephyrin trimer around the J-DNA tip. Second, a correlation plot between 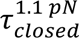 and the hit rate *Q*, that is the likelihood at which an interaction is detected at any given pulling-cycle, showed that around a quarter of the complexes reside in a state where the lifetime is roughly three-fold shorter than usual and in which interactions are seen only one third as often (**Supplementary Fig. S13**). Third, in very rare cases, gephyrin and GlyR β-loops appeared to be trapped in a bound state for extended durations. Approximately 0.05% of all 22577 interactions from a total of N = 94 scaffolds observed during force-scan measurements did not dissociate for at least 15 pulling cycles, and 0.01% of the 84894 interactions (N = 49 scaffolds) observed during constant-force measurements lasted longer than 3400 s (see **Methods** and **Supplementary Fig. S14**). Although such long-lived binding events might have implications for synapse stability, we excluded these rare events from further analysis, as we currently lack a mechanism to explain them and because their infrequent occurrence precludes statistical analysis.

## Discussion

Together, our data provide conclusive evidence that GlyRs and GABA_A_Rs compete for gephyrin binding sites in the post-synaptic density of inhibitory synapses. It has long been thought that such a competition exists based on the fact that the two receptor types bind to overlapping sites of gephyrin ^12, 13^ and that the receptors can be displaced by interfering molecules targeting the same binding pocket (^36^ and references therein). What is new about the current study is that direct competition occurs between GABA_A_Rs and GlyRs at near-endogenous expression levels in mixed inhibitory synapses, and that this competition depends on the thermodynamic properties and the number of interacting molecules. As such, our study sets the stage for an exact mechanistic understanding of inhibitory synaptic plasticity in living neurons.

### Dynamics underlying receptor competition at inhibitory synapses

Our fluorescence microscopy experiments indicate that the accumulation of inhibitory receptor complexes at synapses (**Fig. 1**) correlates with the relative affinity of the receptors for gephyrin that we measured in single molecule binding experiments using magnetic tweezers (**Fig. 4**). Bi-directional changes in receptor occupancy at synapses were observed for GABA_A_R α3-subunits with increased (α3^N369S^) and decreased binding affinity (α3^F368A/I370A^). Our *in vitro* measurements further substantiate the underlying mechanism of this competition, whereby the lifetime of the interaction is the decisive parameter setting the dissociation equilibrium constant and therefore the ratio between bound and unbound GlyRs and GABA_A_Rs. The use of a molecular scaffold ensures that only interactions between a single gephyrin and a single cytoplasmic loop are measured, and that no higher-order structures are present as can be the case in solution-based measurements such as isothermal titration calorimetry (ITC) or surface plasmon resonance. In a cellular context, the competition of α3^N369S^-containing GABA_A_Rs for gephyrin binding sites is seen as a reduction in the effective binding potential of endogenous GlyRs at inhibitory synapses (**Fig. 2f**).

In line with earlier studies ^12, 13^, we observed a pronounced difference in the affinity of wild-type GABA_A_R and GlyR peptides for gephyrin, the latter having an over ten-fold higher *k_D_* (**Table 1**). It is then the more surprising that GABA_A_Rs can compete efficiently in neurons, as shown by the fact that receptors containing the wild-type GABA_A_R α3-subunits are enriched at mixed inhibitory synapses compared with the binding-deficient variant (**Fig. 1**). This seemingly contradictory finding can be reconciled if we consider the relative expression levels of the two receptor types. Although we do not know the absolute receptor copy numbers, the GABA_A_R levels are clearly sufficient to ensure synaptic accumulation of the receptor despite the presence of tightly bound GlyR complexes. It should be noted that both GlyRs and gephyrin are expressed at endogenous levels in these experiments, whereas recombinant GABA_A_R α3-subunits are transduced using a lentivirus expression system. Since membrane targeting of α3-containing receptors relies on the assembly of heteropentameric GABA_A_R complexes with endogenous subunits ^23^, we can assume that the degree of overexpression of GABA_A_R α3 is tolerable. Indeed, GABA_A_R α3 expression did not alter GlyR levels compared to uninfected neurons (**Supplementary Fig. S3**). The presence of several GABA_A_R subunits in a single pentameric receptor complex capable of binding gephyrin could also strengthen the interaction. Based on these considerations, we conclude that the endogenous expression levels of the competing GlyRs and GABA_A_Rs are commensurate with their respective affinities for gephyrin to ensure an effective competition for synaptic binding sites (discussed in ^36^).

A likely consequence of the different binding affinities of the receptors is that GlyRs are predominantly clustered at synapses, whereas the GABA_A_Rs are partially located in the extrasynaptic membrane. If so, a number of conclusions can be drawn. Firstly, due to their lower residency times the plasticity of GABA_A_Rs should be greater than that of the more stably bound GlyRs, suggesting that weaker interactions are better suited for regulatory processes (see below). Secondly, the fact that clustered GlyRs at synapses are, as it were, taken out of the equation implies that the concentration of extrasynaptic GABA_A_Rs may exceed the free GlyRs, which puts the GABA_A_Rs in a better position to compete for the remaining binding sites. Furthermore, it can be hypothesised that GABA_A_Rs preferentially occupy peripheral sites of the synaptic scaffold, whereas GlyRs should predominate in the more stable regions of the post-synaptic domain. The differential distribution of GABA_A_Rs and GlyRs in separate sub-synaptic domains observed in dual colour SMLM experiments ^21^ supports this hypothesis, even though their distribution may not necessarily be reflected in a clear spatial pattern.

### The GlyR-gephyrin binding affinity revisited

Our single molecule measurements revealed that the GlyR-gephyrin interaction has a *k_D_* in the nanomolar range, confirming earlier studies using ITC that also reported a high affinity of the GlyR β-loop for gephyrin. In absolute terms, however, the published values differ substantially, ranging from 16.5 nM (this study) to low micromolar concentrations (discussed in ^18^). This discrepancy can be partially explained by the length of the β-loop sequence used. Shorter peptide constructs tend to have lower affinity, as seen in our competition experiments with 11mers of the GlyR β-loop (*K_D_*= 2.23 µM). The underlying reason is that the flanking regions of the core sequence contribute to the GlyR binding to gephyrin ^32, 37, 38^.

Several studies have also suggested that the GlyR β-gephyrin interaction is bimodal (e.g. ^16, 39^), with a second, low affinity binding site accounting for up to two-thirds of the interactions. These findings appear to be related, given that bimodal binding in other studies was generally observed when longer β-loop sequences were used ^18,38^. The repeated measurement of single molecule interactions using magnetic tweezers can provide an independent assessment of the GlyRβ-gephyrin binding model. Although we observed bimodal interactions on rare occasion, these were not the rule. The distribution of dwell times of the closed conformation followed a single exponential (**Fig. 3b and Fig. 5c**). A second binding site with a higher dissociation rate constant would have manifested itself by a second decay, as long as the binding events can be detected by our setup.

When looking for changes in the extension amplitude, however, we did detect around 20% of scaffolds for which the Δ*z* histogram was bimodal (**Supplementary Fig. S12e**). This phenomenon is likely due to the sterically flexible arrangement of trimeric gephyrin, in particular its unstructured C-domain, and to the fact that the β-loop can interact with more than one of the gephyrin subunits (e.g. ^35^). In other words, the observed bimodality of the extension jump does not correspond to the two-site model described above, since in our experiments the lifetimes of the different extension states were not altered.

### Single molecule biochemical measurements *in vitro* and *in cellula*

Recent studies have described the formation of synaptic gephyrin clusters in terms of phase separation, where condensation is driven by various intra- and intermolecular interactions between inhibitory receptors and scaffold proteins ^38, 40^. Here, we demonstrate that receptor-gephyrin diffusion and trapping at synapses depend critically on specific interactions between individual molecules, and that these interactions can be accurately measured and quantified in the absence of aggregation. After all, our magnetic tweezer measurements with J-DNA constructs ensure the interaction of single pairs of molecular partners, contrary to bulk techniques for which a doubt always remains whether higher-order structures could form. By relating the *in vitro* binding data with single particle tracking (SPT) of receptor diffusion, we quantitatively describe the dynamic equilibrium at inhibitory synapses in a physiological (cellular) context. The comparability of the two datasets is ensured by the use of single molecule approaches that can account for temporal changes in the binding behaviour, resulting for instance from post-translational modifications, changes in conformation and/or binding geometry of the different components.

Our J-DNA measurements of receptor-gephyrin interactions revealed large differences in the interaction lifetimes of the intracellular receptor loops and gephyrin. From the dissociation equilibrium constants we can further deduce energetic parameters of the receptor-gephyrin interactions, characterizing them by their free energy of binding according to Δ*G* = *k_B_T* ln *k_D_*. Comparing the results with the different peptide sequences, we obtain a binding free energy of Δ*G_β_* = −13.1 ± 1 *k_B_T* for the GlyR β-peptide-gephyrin interaction, which is substantially stronger than the GABA_A_R α3^WT^-gephyrin binding free energy of Δ*G_α_*_3_ = −10.1 ± 1.6 *k_B_T*. The GABA_A_R α3^N369S^-gephyrin interaction displays an intermediate binding free energy of Δ*G_α_*_3_ *_N_*_369*S*_ = −11.7 ± 1.6 *k_B_T*.

The free energies of receptor-gephyrin binding are consistent with the trapping of the receptors at synapses. Using PALM-based SPT recordings in live neurons, we derived an energy landscape that measures the depth of the potential well of the post-synaptic gephyrin network with which the diffusing GlyRs interact. Bi-directional changes in potential energy were observed in the presence of competing receptor variants that differ in binding free energy. An important difference between the SPT diffusion data and the molecular tweezer data is that SPT measures the mobility of individual receptors in the presence of densely packed protein clusters. In other words, the mobility reflects the effective diffusion of the membrane receptors undergoing rapid binding and unbinding at neighbouring sites. Interestingly, our diffusion data indicates that the mobility (or diffusivity) of the GlyR is not altered in the presence of competing receptors, whereas its effective trapping energy at synapses is reduced. This observation fits well with the notion that the lifetime of the receptor-scaffold interactions *τ*_closed_ is a key parameter regulating the level of gephyrin binding-competent receptor complexes at synapses rather than the receptor diffusion coefficients.

### A role for receptor competition in synaptic plasticity

What could be the purpose of a common binding site, when one of the binding partners can be more easily displaced than the other? The coexistence of inhibitory receptor types with different affinity for the post-synaptic gephyrin scaffold at mixed synapses could reconcile the need of long-term synapse stability with the continuous exchange of its components and plastic remodelling on a fast timescale. Our magnetic tweezer data show that GABA_A_Rs are much less strongly bound to gephyrin than GlyRs. If GABA_A_Rs needed tighter binding, a simple point mutation could easily achieve this, as shown by the α3^N369S^ variant. The fact that the wild-type GABA_A_R α3-subunit does not contain such a mutation suggests that the affinity of the receptor is well-adjusted to a physiological function that requires a greater level of mobility ^41^, including a role in tonic inhibition (e.g. ^42^) or alternative, gephyrin-independent clustering mechanisms ^43, 44^. As an example of the greater plasticity of GABA_A_Rs compared to GlyRs, it was shown that cAMP-dependent phosphorylation of gephyrin at residue S270 reduces GABA_A_R binding at synapses and increases receptor diffusion, whereas GlyRs are largely unaffected ^45^.

At the same time, mixed inhibitory synapses in the spinal cord require a high degree of stability to ensure reliable signal transmission in sensory and motor circuits ^8^. The binding of high-affinity GlyRs to synaptic gephyrin clusters leads to high receptor occupancy that is estimated at about 50% ^19^. Due to their lower affinity, GABA_A_ receptors at these synapses possibly play only a modulatory role, for example to provide functional compensation in a pathological setting. It should be noted that the number of receptor binding sites at inhibitory synapses is by no means constant. The stereotypical packing of GlyRs at spinal cord synapses argues for a modular arrangement, in which the addition or removal of receptor-gephyrin complexes regulates the size of the inhibitory post-synaptic domain ^19^. In other words, the loss of receptors is invariably accompanied by a (partial) dispersal of the synaptic gephyrin scaffold. In our present study, the expression of the high affinity GABA_A_R α3^N369S^ variant increases the size of the post-synaptic scaffold (**Fig. 1c**). The efficacy of different inhibitory receptors to compensate the loss of GlyRs in pathologies such as startle disease or hyperekplexia is therefore likely to depend not only on their ability to compete for synaptic binding sites, but also their capacity to maintain and stabilise these sites ^20^.

In conclusion, it can be said that two key parameters define the equilibrium between competing membrane proteins at synapses: the lifetime of the interactions and the concentration of the different binding partners. This is the case for dynamic equilibria in general and should therefore apply to all types of competition, including the replacement of like with like, for instance the exchange of desensitised AMPA receptors that was shown to underlie the recovery from synaptic depression at glutamatergic synapses ^46^. Knowledge of these two parameters is therefore essential for biophysical modelling that can provide a fully quantitative description of a given molecular system in living cells.

## Methods

### In-cell experiments and fluorescence microscopy

#### Primary spinal cord neuron culture

All experiments were in accordance with the European Union guidelines and approved by the local veterinary authorities. Animals at IBENS were treated in accordance with the guidelines of the French Ministry of Agriculture and Direction Départementale des Services Vétérinaires de Paris (École Normale Supérieure, Animalerie des Rongeurs, license B 75-05-20).

Primary spinal cord neuron cultures were prepared at embryonic day 13.5 (E13.5) from homozygous mEos4b-GlyRβ knock-in (KI) mice where the β-subunit of the GlyR is tagged with the mEos4b fluorescent protein at the N-terminal extracellular site (*Glrb*^Eos/Eos^; mouse strain C57BL/6N-*Glrb^tm1Ics^*, backcrossed into C57BL/6J, accession number MGI:6331106, ^19^). Spinal cord tissue was treated with the protease papain for 10 min at 37°C, triturated in BSA in the presence of DNase I (final concentration 0.1 mg/ml), and centrifuged at 1000 rpm for 8 min with a BSA cushion (2 mg/mL). Neurons were plated at a density of 2.5 × 10^5^ cells/cm^2^ onto 18 mm diameter glass coverslips, #1.5 thickness (VWR), precoated with 80 µg/ml poly-L-ornithine (Sigma). Neurons were kept at 37°C and 5% CO_2_ in neurobasal medium (Thermo Fisher Scientific) containing 2 mM L-glutamine (Thermo Fisher Scientific), B27 (1x) supplement (Thermo Fisher Scientific) and penicilin/streptomycin (P/S; Thermo Fisher Scientific) with an exchange of half the medium every 4 days in culture.

#### Expression constructs and lentivirus infection

Lentivirus constructs for the expression of recombinant GABA_A_R α3-subunits were cloned in the FUGW backbone ^47^, containing a generic signal peptide (SP) derived from the human *GLRB* gene followed by the coding sequence (CDS) of mScarlet, and the full-length coding sequence (exluding the SP) of rat *Gbra3* (gift from U. Zeilhofer). This FU-mScar-rGabra3^WT^ construct was modified by site-directed mutagenesis to generate FU-mScar-rGabra3^N369S^ and FU-mScar-rGabra3^F368/I379A^. The corresponding Halo-tagged constructs were cloned in the same way in an FUGW backbone containing the SP of rat *Gbrg2*, a myc and a Halo-7 tag, followed by the CDS of wild-type or mutated rat *Gbra3*, giving rise to the constructs FU-Halo-rGabra3^WT^, FU-Halo-rGabra3^N369S^ and FU-Halo-rGabra3^F368/I379A^.

Lentiviruses were produced in HEK-293 cells as previously described ^45^. Briefly, cells were co-transfected with FUGW replicon DNA and the third generation helper plasmids pMD2.G, pMDLg/pRRE, and pRSV-Rev (Addgene #12259, #12251, #12253) with lipofectamine 2000 (Thermo Fisher Scientific) according to supplier’s instructions. Cells were maintained in Neurobasal medium containing GlutaMAX, B27 and P/S at 32°C / 5% CO_2_. The medium was replaced at 24 h after transfection, and the lentivirus-containing medium was collected at 48 - 55 h, cleared by filtration (0.45 μm pore size), and stored at −80°C. Neurons were infected at DIV4 by adding the lentivirus directly to the medium.

#### Immunolabelling and epifluorescence imaging

Epifluorescence imaging in fixed samples was carried out in FU-mScar-rGabra3 infected neurons. Neurons were fixed at DIV18 for 20 min at room temperature in 4% (w/v) paraformaldehyde (Electron Microscopy Sciences) and 4% sucrose (w/v) in phosphate buffered saline (PBS, pH 7.4), followed by 3 washes in PBS. Neurons were blocked in 3% (w/v) bovine serum albumin (BSA, Sigma Aldrich) and 0.25% triton-X100 (Sigma) in PBS for 1 h at room temperature, followed by incubation with the primary mouse monoclonal anti-gephyrin antibody mAb7a (Synaptic Systems #147011; 1:500) for 1 h in fresh blocking buffer at room temperature. Neurons were washed 3 times in PBS and incubated with the secondary antibody donkey anti-mouse AF647 (Invitrogen; 1:1000) in fresh blocking buffer for 1 h at room temperature, followed by 3 washes in PBS.

Imaging was done on an inverted Nikon Eclipse Ti microscope with a 100x/1.49 NA oil-immersion objective and an additional 1.5x magnifying lens, using an Andor iXon EMCCD camera (16-Bit, 107 nm final pixel size), and NIS-Elements software (Nikon). Ten images of 512 x 512 pixels with an exposure time of 100 ms (for mEos4b and mScarlet) or 200 ms (for AF647) per frame were taken with lamp illumination, using specific filters for mEos4b (excitation: 485 nm, emission: 525 nm), mScarlet (excitation: 560, emission: 607) and AF647 (excitation: 650, emission: 684).

#### Epifluorescence image analysis

For all images of mEos4b-GlyR β, mScarlet-GABA_A_R α3 and gephyrin-7a-AF647, an average projection was generated from the 10 frames captured for each channel. In order to analyse the intensity of mEos4b-GlyR β and mScarlet-GABA_A_R α3 at inhibitory synapses in spinal cord neurons, we first ran the ICY Spot Detector plug-in ^48^ on the gephyrin-7a-AF647 images, to produce a set of regions of interest (ROI). Using the image analysis software Fiji, these ROIs were then used to measure the raw integrated intensities of mEos4b-GlyR β, mScarlet-GABA_A_R α3 and gephyrin-7a-AF647 at each synaptic gephyrin clusters.

#### Single particle tracking (SPT)

SPT imaging based on live PALM recordings (sptPALM) of endogenous mEos4b-GlyR β was carried out in FU-Halo-rGabra3 infected neurons at DIV18-20. Coverslips were transferred to a Ludin chamber with artificial cerebral spinal fluid (ACSF) imaging solution (127 mM NaCl, 3 mM KCl, 2 mM CaCl_2_, 1.3 mM MgCl_2_, 10 mM glucose, 10 mM HEPES, pH adjusted to 7.38), pre-warmed to 37°C. JF646-Halo-ligand (Promega) was added at a final concentration of 20 nM along with 0.5 μL 100 nm-sized Tetraspeck beads (Invitrogen) for 15 min at 37°C. Neurons were then washed twice and imaged in 1 mL imaging solution. SPT imaging was carried out on an inverted Nikon Eclipse Ti microscope with a ×100/1.49 NA oil-immersion objective and an additional 1.5x lens using an Andor iXon EMCCD camera (16-Bit, 107 nm pixel size), and NIS-Elements software (Nikon). The imaging chamber was heated to 35°C. Images of 200 x 200 pixels were acquired. Lamp images were first taken of the unconverted mEos4b-GlyR β (10 frames of 100 ms), followed by movies of 20 frames of JF-646/Halo-tagged GABA_A_R α3-subunits with continuous 633 nm laser illumination (nominal laser power 400 mW, output set at 20%, 100 ms frames) to identify lentivirus infected neurons. Movies of 40000 frames were then recorded using a red 561 nm imaging laser operated in continuous mode (nominal laser power 200 mW, set at 50%, 15 ms frames, emission filter 607/36). Photoconversion of mEos4b-GlyR β was done through 0.5 ms pulsed illumination with a 405 nm laser (output power set at 8%) together with a 488 nm laser (nominal laser power 100 mW, output 10%) to increase the length of the SPT tracks of mEos4b as described in ^49^. The focal plane was maintained using a Nikon perfect focus system.

#### PALM and SPT analysis

Quantification of mEos4b-GlyR β was carried out using a lab script for MATLAB. The mEos4b single fluorophores were detected by Gaussian fitting. The resulting pointillist images were drift corrected using the Tetraspeck beads identified in the projected pointillist PALM images. Rendered images were produced with a pixel size of 10 nm and σ = 10 nm. A lab written script for MATLAB was used to reconstruct mEos4b-GlyR β trajectories. The Python script TRamWAy ^26^ was then used to perform spatio-temporal analysis to map the effective diffusivity and effective energy of mEos4b-GlyR β at synapses.

#### Statistical analyses

Graphing and statistical analyses of the imaging data shown in **Fig. 1, 2 and Supplementary Fig. S1-S3** were carried out using Python3. Data were tested for normality of distribution using a D’Agostino Pearson test. *p<0.05, **p<0.01, ***p<0.001, ns (not significant). For the remaining figures the Xvin software suite and statistical analysis packages were used (PicoTwist SARL).

### In vitro magnetic tweezers measurements

#### J-DNA synthesis

Comprehensive descriptions of the nucleic acid scaffolds are given in previous publications ^27, 33, 50^. Briefly, J-DNA forceps consist of two linear dsDNA segments or branches, connected via a third segment, the leash. The juncture with the leash divides each branch into a tip and a shank. The proteins are engrafted at the end of the tips; the shanks are ligated to ≈ 1000 bp-long dsDNA fragments that are multiply labelled with either biotin or digoxigenin and used for the attachment to a magnetic bead or to the bottom window of a flow cell. The scaffold is symmetric, both tips being 48 bp-long and both shanks being ≈ 1500 bp-long, which clearly differentiates specific interactions between the molecular partners from non-specific interactions between one of the proteins and the surfaces ^34^ (**Supplementary Table 1**). The leash is ≈ 700 bp long to maximize the chances of association of the complex ^27^. For grafting we used specific, stable thioether bonds between a small molecule located at each J-DNA tip, either benzylguanine or benzylcytosine, and a protein tag located at the N-terminus of the two proteins, either SNAP or CLIP, respectively ^27, 51, 52^ (see **Supplementary Information** for details).

#### Protein production

*In vitro* experiments were performed with the full-length gephyrin rC4 splice variant that was fused at its N-terminus with a 6xHis-SNAP-tag (**Supplementary Fig. S4**), expressed in *E. coli*, and purified as described previously ^32^. The purified 6xHis-SNAP-gephyrin construct formed trimers as confirmed by chromatography (**Supplementary Information and Fig. S5**). As regards the receptor loops, the GlyR β, GABA_A_R α3^WT^ and GABA_A_R α3^N369S^ sequences were inserted inside the pleckstrin homology (PH) domain of human cytohesin-1 (**Supplementary Fig. S4**) as described in ^53^, and fused at the N-terminus of the PH-domain with a CLIP-tag for attachment to the J-DNA tips (**Supplementary Fig. S4**). All three constructs (10xHis-CLIP-GlyR β-, 10xHis-CLIP-GABA_A_R α3^WT^-, and 10xHis-CLIP-GABA_A_R α3^N369S^-loops) were expressed in *E. coli* and purified by affinity chromatography, cation exchange, and gel filtration (see **Supplementary Information** for details).

#### Peptide synthesis

The GlyR β-peptide (sequence FSIVGSLPRDF, MW = 1237.42 g/mol), GABA_A_R α3^WT^-peptide (FNIVGTTYPINC, 1341.54 g/mol), and GABA_A_R α3^N369S^-peptide (FSIVGTTYPINC, 1314.52 g/mol) were synthesized as previously described ^54^. Single-use aliquots of stock solutions in water (2 mM for the GlyR β-peptide and 10 mM for GABA_A_R α3^WT^ and GABA_A_R α3^N369S^) were stored at −80°C.

#### Setup operation

Measurements were carried out on homemade magnetic tweezers as described previously ^28, 55, 56^ (see also **Supporting Information**). The setup was built on an inverted microscope equipped with a video acquisition system that monitors the position of the magnetic beads in 3D (with 5 nm precision) and in real-time (at up to 90 Hz). Magnets on a translation stage above the sample are used to vary the force applied to the nucleic acid scaffold from 0 to 10 pN, with a ≈ 10% precision ^27, 28, 57, 58^. Peltier devices affixed to the oil-immersion objective set the temperature of the J-DNA from 19 to 35°C within ± 0.5°C.

Nanomanipulation is done in a flow cell made of two glass coverslips, the upper with two access holes connecting to reservoirs for buffer exchange. The two windows are separated by a double layer of parafilm with a central slit that delineates a fluidic channel ^28, 59^. Two surface functionalization methods were used to attach the J-DNAs to the lower coverslip: either the glass was coated with a thin layer of polystyrene for subsequent non-specific absorption of anti-digoxigenin ^27, 28, 59, 60^, or the glass was PEGylated with some of the polymer molecules containing a biotin group at one end for streptavidin binding ^59^. Titrations at constant force were done using the second approach, whereas both methods were applied for the cycling-force experiments. All measurements were done in gephyrin buffer (20 mM Tris pH 7.5, 250 mM NaCl, 5 mM MgCl_2_, 0.1% Tween20, 0.5 mg/mL BSA, 5 mM β-mercaptoethanol). The temperature was set at 19.2°C unless stated otherwise.

#### Sample preparation

Depending on the used proteins, the corresponding functionalization was selected for the magnetic beads: either we used commercial particles coated with streptavidin (MyOne Streptavidin C1, Thermo Fisher Scientific), or we coated with anti-digoxigenin particles displaying reactive groups at their surface (MyOne Tosylactivated, Thermo Fisher Scientific)^59^. In both cases the 1 μm-diameter beads were mixed with J-DNAs and the resulting assemblies injected in the fluidic channel, allowing their binding to the bottom window until the field of view displayed 50 to 70 tethered objects ^27, 28, 33^. Unbound particles were rinsed out. Next, SNAP and CLIP-tagged proteins were injected at 100 to 500 nM in gephyrin buffer, either sequentially or together, and incubated with the DNA scaffolds for 2 to 15 h at 19.2°C. The unbound proteins were washed out from the flow chamber with 10 to 20 mL of buffer before measurements could begin. Typically, in a single field-of-view up to 30% of the J-DNAs displayed interactions, in agreement with previous experiments on other biological systems ^27, 28^.

#### Constant force experiments and data analysis

Measurements were performed in gephyrin buffer and at 19.2°C, using a gephyrin trimer and the GlyR β-loop engrafted at the tips of J-DNAs, based on earlier protocols ^28, 29^. The force was kept constant at 40 fN, a necessary compromise to distinguish the closed and open conformations while maintaining the proximity of the dissociated partners allowing them to re-associate quickly. The titrant was added in solution at 5 to 7 different concentrations between 0 and ≈ 1.5 *k_D_* with increasing concentration. Above 1 *K_D_* the number of closing transitions is very low, demanding excessively long acquisitions (**Supplementary Fig. S7b, d, and f**). In practice, data were collected for 7.5 to 23 h, the goal being to observe at least ≈ 100 closing-opening cycles per scaffold. The reversibility of the titration was shown in a separate recording (**Supplementary Fig. S15**).

Time traces were automatically analysed with a Python program based on hidden Markov modelling ^28^. After identification of both closed and open conformations, transitions were detected and dwell times determined. Dwell times Δ*t_closed_* and Δ*t_open_* are shown as histograms and fitted by monoexponentials, yielding the characteristic times *τ_closed_* and *τ_open_*. This analysis is rigorous in the first case, whereas in the presence of titrant we only approximate the true Δ*t_open_* distribution by its slowest decay ^28, 29^. Also, assimilating 1/*τ_closed_* at 40 fN to the dissociation rate constant in solution, i.e. at 0 fN, is fully legitimate when considering the ≈ 7 Å Bell parameter for this interaction (**Supplementary Fig. S10**): the error is below 1%. Single-scaffold histograms displaying less than 50 events after removal of the first bin were discarded. Likewise, long events that did not dissociate within 3400 s were excluded. Traces with less than 50 dissociations or associations were not considered for titration (**Supplementary Table S2**). Furthermore, for each titrant only the scaffolds that endured at least 3 concentrations were retained (N = 8 - 10, **Supplementary Table S2**).

Obtaining the bimolecular association rate constant, *k_A_*, and the dissociation equilibrium constant, *k_D_*, was more complicated, since the binding steps we detect with J-DNAs are monomolecular events and their properties strongly depend on several experimental parameters such as the length of the leash connecting the two partners and the applied force. *k_D_* was determined for each individual scaffold using the mass action law expressed in terms of fractional occupancies:

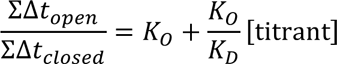

with *k_D_* the *in trans* complex dissociation equilibrium constant, *k_O_*= *k_D_* 3*C_eff_* the scaffold opening equilibrium constant, and *C_eTT_* the effective concentration at the tips ^28, 29^. After normalization by the value measured in absence of titrant we obtain:

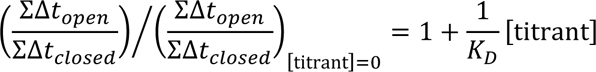

showing that linear fits are a straightforward method to extract *k_D_*, assuming no cooperativity between the different binding sites of the gephyrin trimer. For each competitor the results were averaged over all the scaffolds and the distribution mean, < *k_D_* > and error were extracted.

The association rate constant for GlyRβ-loop (but not the other competitors), *k_A_*, was first derived for each individual scaffold, as 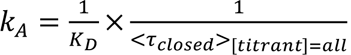 with < *τ_closed_* >_[_*_titrant_*_]=*all*_ corresponding to the averaged value over all concentrations of the competitor. Subsequently, the results were averaged over all the scaffolds and the distribution mean, < *k_A_* >, was calculated and the error propagated.

#### Cycling force experiments and data analysis

Measurements were done with a gephyrin trimer and either GlyR β, GABA_A_R α3^WT^, or GABA_A_R α3^N369S^-loops engrafted at the tips of J-DNAs, following established protocols ^27, 29,33^. In all cases a low force of 0.001 pN was applied for 12 - 15 s. The temperature was generally fixed at 19.2°C and a high force of 1.1 pN was typically applied for a duration of 155 s for the GlyR β-loop, 1 s for GABA_A_R α3^WT^-loop, and 10 s for the N369S variant. The only exceptions to these setting are the recordings of the gephyrin-GlyR β-loop energy landscape, in which the temperature ranged from 19.2 to 24.1 °C and the high force from 1.1 to 9.6 pN (**Supplementary Fig. S10**). Tension was applied in this case for 55 - 190 s, shorter durations being used at higher force. Data were collected on 5 to 50 individual scaffolds for 3.5 - 114 h, collecting between 63 and 772 of individual events per scaffold.

Typically, the measurements were performed at the 31 Hz acquisition rate and at least three points present at the expected intermediate extension on the time trace (corresponding to a 97 ms detection limit) were required to confirm that an event took place (**Fig. 5b**). A faster camera with a 90 Hz acquisition rate was used for GABA_A_R α3^WT^-loop (see **Supplementary Information**), to measure dwell times as short as 55 ms (five data points).

Results were analysed using Xvin software (PicoTwist) as in ^27, 33^. Histograms of the dwell times Δ*t*_*closed*_ were built and processed as in the constant force experiments. Single-scaffold histograms displaying less than 50 events before removal of the first bin were discarded. Long events, for which unbinding did not occur within the same cycle as the one in which binding was first observed were excluded.

For GlyR β-loop-gephyrin complex, individual events from each scaffold were fit to single-exponential distribution, and subsequently either fit to Gaussian distribution (data from N = 50 scaffolds collected at F = 1.1 pN and T = 19.2 °C, see **Supplementary Figure S13**) or averaged (all remaining conditions, data from N = 5 – 10 scaffold). From data collected at F = 1.1 pN and T = 19.2 °C, for each scaffold, we also evaluated the hit rate *Q*, defined as the ratio between the number of detected rupture events upon the total number of force cycles contained in a time trace (see **Supplementary Figure S13**). GABA_A_R α3^WT^-loop-gephyrin complex was very short-lived, and the events with a dwell time below 55 ms, were severely undercounted. We also noted that GABA_A_R α3^N369S^-loop-gephyrin complex was not observed as frequently as when GlyR β-loop was engrafted on the scaffold. Additionally, tip-specific interactions appeared infrequently and at times clustered. We were not able to determine factors responsible for this behavioural heterogeneity but possibly it can be related to different than for GlyR β-loop dynamics of the nested GABA_A_R α3^N369S^-loop within the PH-domain.

Amplitude changes upon rupture, Δ*z*, were represented as histograms, error bars corresponding to counting errors and fit to single or double Gaussian distributions. Typically, the expected Δ*z* signature for specific interactions lies between 130 and 200 nm, depending on the applied force (see **Supplementary Table S1** for a computation based on the contour length). All events that were not within three standard deviations of the central value were not taken into account.

To map the energy landscape of the GlyR β-loop-gephyrin complex we measured *τ_closed_* at different force and temperature. Results collected with individual scaffolds were averaged and standard errors were calculated. The logarithm of the dissociation rate constant under force *F* was then determined as − ln*τ_closed_* and errors propagated. Finally, a weighted, least squares fit to the linearized Bell equation:

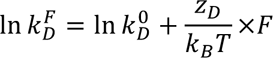

provided the dissociation rate constant at zero-force, 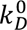, and the distance to the transition *zD*, and the lifetime of interaction at zero-force, *τD0* (as 1/*kD0*) (**Supplementary Fig. S10a**). The activation energy, *E_D_*, was obtained by fitting data collected at 1.1 pN to the linearized Arrhenius equation:

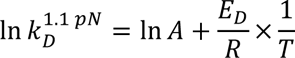

(**Supplementary Fig. S10b**).

#### Analysis of rare, long-lived interactions

During constant-force measurements, we identified 7 out of 84894 analysed closed events characterized by dwell times (Δ*t_closed_*) ranging from 3749 s to over 76859 s (Δ*t_closed_* of some interactions exceeded the duration of the experiment, and therefore are indeterminate). The analysis employed a cut-off of 3400 s, representing a 150-fold difference from the mean interaction lifetime (<*τ_closed_*> = 22.7 ± 0.4 s) observed under the same experimental conditions. These long-lived events were detected in 7 out of the 49 analysed individual scaffolds.

During cycle-force measurements, we identified 11 out of 22577 analysed closed events that remained intact for at least 15 consecutive pulling cycles without dissociation. The duration of these closed events ranged from 16 to over 405 cycles. In some instances, the interactions did not dissociate within the duration of the entire measurement, making the exact number of cycles indeterminate. Similar to the constant-force measurements, we set a cut-off of 15 pulling cycles corresponding to an approximately 150-fold time difference from the mean interaction lifetime (<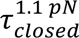 > = 16.9 ± 0.8 s) observed under a force of F = 1.1 pN at 19.2 °C. These long-lived events were detected for 11 out of 94 analysed individual scaffolds.

## Acknowledgements

We thank Uli Zeilhofer (University of Zürich) and Theofilos Papadopoulos (MPI Göttingen) for the original Gabra3 plasmid and its modification, Nora Grünewald and Günter Schwarz (Universität Köln) for the gephyrin expression constructs and protocols, and François Stransky (IBENS Paris) for the analysis pipeline of the single-molecule titration data. Our research received funding from Labex Memolife (project Gephyrip to TRS and CGS) and ANR (project InVivoNanoSpin to CGS). SAM was supported by a Fondation pour la Recherche Médicale (FRM) postdoctoral fellowship (SPF201809007132).

## Author contributions

DK and SAM performed the experiments and analysed the data; CS, FL, JBM, HMM, CG, and CGS developed and characterised experimental and analytical tools; DK, SAM, CG, and CGS wrote the manuscript with contributions of AT and TRS; all authors participated in critical reading and editing of the article.

## References

1. O’Brien JA, Berger AJ. Cotransmission of GABA and glycine to brain stem motoneurons. Journal of neurophysiology, (1999).

2. Dufour A, Tell F, Kessler JP, Baude A. Mixed GABA–glycine synapses delineate a specific topography in the nucleus tractus solitarii of adult rat. The Journal of physiology 588, 1097–1115 (2010).

3. Nabekura J, et al. Developmental switch from GABA to glycine release in single central synaptic terminals. Nature neuroscience 7, 17–23 (2004).

4. Dumoulin A, Triller A, Dieudonne S. IPSC kinetics at identified GABAergic and mixed GABAergic and glycinergic synapses onto cerebellar Golgi cells. J Neurosci 21, 6045–6057 (2001).

5. Jonas P, Bischofberger J, Sandkuhler J. Corelease of two fast neurotransmitters at a central synapse. Science 281, 419–424 (1998).

6. Tanaka I, Ezure K. Overall distribution of GLYT2 mRNA-containing versus GAD67 mRNA-containing neurons and colocalization of both mRNAs in midbrain, pons, and cerebellum in rats. Neuroscience research 49, 165–178 (2004).

7. Bohlhalter S, Mohler H, Fritschy J-M. Inhibitory neurotransmission in rat spinal cord: co-localization of glycine-and GABAA-receptors at GABAergic synaptic contacts demonstrated by triple immunofluorescence staining. Brain research 642, 59–69 (1994).

8. Alvarez FJ. Gephyrin and the regulation of synaptic strength and dynamics at glycinergic inhibitory synapses. Brain Res Bull 129, 50–65 (2017).

9. Bardoni R, Takazawa T, Tong CK, Choudhury P, Scherrer G, MacDermott AB. Pre-and postsynaptic inhibitory control in the spinal cord dorsal horn. Annals of the New York Academy of Sciences 1279, 90–96 (2013).

10. Dumoulin A, Levi S, Riveau B, Gasnier B, Triller A. Formation of mixed glycine and GABAergic synapses in cultured spinal cord neurons. Eur J Neurosci 12, 3883–3892 (2000).

11. Maynard SA, Ranft J, Triller A. Quantifying postsynaptic receptor dynamics: insights into synaptic function. Nature Reviews Neuroscience 24, 4–22 (2023).

12. Maric HM, Mukherjee J, Tretter V, Moss SJ, Schindelin H. Gephyrin-mediated gamma-aminobutyric acid type A and glycine receptor clustering relies on a common binding site. J Biol Chem 286, 42105–42114 (2011).

13. Maric HM, et al. Molecular basis of the alternative recruitment of GABA(A) versus glycine receptors through gephyrin. Nat Commun 5, 5767 (2014).

14. Maric HM, Kasaragod VB, Schindelin H. Modulation of gephyrin-glycine receptor affinity by multivalency. ACS Chem Biol 9, 2554–2562 (2014).

15. Mukherjee J, et al. The residence time of GABA(A)Rs at inhibitory synapses is determined by direct binding of the receptor alpha1 subunit to gephyrin. J Neurosci 31, 14677–14687 (2011).

16. Specht CG, Grunewald N, Pascual O, Rostgaard N, Schwarz G, Triller A. Regulation of glycine receptor diffusion properties and gephyrin interactions by protein kinase C. EMBO J 30, 3842–3853 (2011).

17. Tretter V, et al. Molecular basis of the gamma-aminobutyric acid A receptor alpha3 subunit interaction with the clustering protein gephyrin. J Biol Chem 286, 37702–37711 (2011).

18. Kasaragod VB, Schindelin H. Structure-function relationships of glycine and GABAA receptors and their interplay with the scaffolding protein gephyrin. Front Mol Neurosci 11, 317 (2018).

19. Maynard SA, et al. Identification of a stereotypic molecular arrangement of endogenous glycine receptors at spinal cord synapses. Elife 10, e74441 (2021).

20. Wiessler AL, et al. Role of the glycine receptor β subunit in synaptic localization and pathogenicity in severe startle disease. J Neurosci 44, (2024).

21. Yang X, Le Corronc H, Legendre P, Triller A, Specht CG. Differential regulation of glycinergic and GABAergic nanocolumns at mixed inhibitory synapses. EMBO Rep 22, e52154 (2021).

22. Bohlhalter S, Weinmann O, Mohler H, Fritschy J-M. Laminar compartmentalization of GABAA-receptor subtypes in the spinal cord: an immunohistochemical study. Journal of Neuroscience 16, 283–297 (1996).

23. Hannan S, Smart TG. Cell surface expression of homomeric GABA(A) receptors depends on single residues in subunit transmembrane domains. J Biol Chem 293, 13427–13439 (2018).

24. Laurent F, Floderer C, Favard C, Muriaux D, Masson JB, Vestergaard CL. Mapping spatio-temporal dynamics of single biomolecules in living cells. Phys Biol 17, 015003 (2019).

25. Masson JB, et al. Mapping the energy and diffusion landscapes of membrane proteins at the cell surface using high-density single-molecule imaging and Bayesian inference: application to the multiscale dynamics of glycine receptors in the neuronal membrane. Biophys J 106, 74–83 (2014).

26. Laurent F, Verdier H, Duval M, Serov A, Vestergaard CL, Masson J-B. TRamWAy: mapping physical properties of individual biomolecule random motion in large-scale single-particle tracking experiments. Bioinformatics 38, 3149–3150 (2022).

27. Kostrz D, et al. A modular DNA scaffold to study protein–protein interactions at single-molecule resolution. Nature nanotechnology 14, 988–993 (2019).

28. Stransky F, et al. Use of DNA forceps to measure receptor-ligand dissociation equilibrium constants in a single-molecule competition assay. Methods in Enzymology 694, 51–82 (2024).

29. Saha P, et al. Modulation of SARS-CoV-2 spike binding to ACE2 through conformational selection. bioRxiv, 2024.2003. 2015.585207 (2024).

30. Han L, et al. Concentration and length dependence of DNA looping in transcriptional regulation. PLoS One 4, e5621 (2009).

31. Rieu M, Valle-Orero J, Ducos B, Allemand JF, Croquette V. Single-molecule kinetic locking allows fluorescence-free quantification of protein/nucleic-acid binding. Communications biology 4, 1083 (2021).

32. Grunewald N, et al. Sequences flanking the gephyrin-binding site of GlyRbeta tune receptor stabilization at synapses. eNeuro 5, ENEURO.0042–0017.2018 (2018).

33. Wang JL, et al. Dissection of DNA double-strand-break repair using novel single-molecule forceps. Nature structural & molecular biology 25, 482–487 (2018).

34. Wang YJ, et al. Combining DNA scaffolds and acoustic force spectroscopy to characterize individual protein bonds. Biophysical Journal 122, 2518–2530 (2023).

35. Sander B, et al. Structural characterization of gephyrin by AFM and SAXS reveals a mixture of compact and extended states. Acta Crystallogr D Biol Crystallogr 69, 2050–2060 (2013).

36. Specht CG. Fractional occupancy of synaptic binding sites and the molecular plasticity of inhibitory synapses. Neuropharmacology 169, 107493 (2020).

37. Maric HM, et al. Design and synthesis of high-affinity dimeric inhibitors targeting the interactions between gephyrin and inhibitory neurotransmitter receptors. Angew Chem Int Ed Engl 54, 490–494 (2015).

38. Bai G, Wang Y, Zhang M. Gephyrin-mediated formation of inhibitory postsynaptic density sheet via phase separation. Cell research 31, 312–325 (2021).

39. Kim EY, et al. Deciphering the structural framework of glycine receptor anchoring by gephyrin. Embo J 25, 1385–1395 (2006).

40. Lee G, et al. Thermodynamic modulation of gephyrin condensation by inhibitory synapse components. Proceedings of the National Academy of Sciences 121, e2313236121 (2024).

41. Petrini EM, et al. Synaptic recruitment of gephyrin regulates surface GABAA receptor dynamics for the expression of inhibitory LTP. Nat Commun 5, 3921 (2014).

42. Chery N, de Koninck Y. Junctional versus extrajunctional glycine and GABA(A) receptor-mediated IPSCs in identified lamina I neurons of the adult rat spinal cord. J Neurosci 19, 7342–7355 (1999).

43. Kneussel M, Brandstatter JH, Gasnier B, Feng G, Sanes JR, Betz H. Gephyrin-independent clustering of postsynaptic GABA(A) receptor subtypes. Mol Cell Neurosci 17, 973–982 (2001).

44. Panzanelli P, et al. Distinct mechanisms regulate GABAA receptor and gephyrin clustering at perisomatic and axo-axonic synapses on CA1 pyramidal cells. J Physiol 589, 4959–4980 (2011).

45. Niwa F, Patrizio A, Triller A, Specht CG. cAMP-EPAC-Dependent Regulation of Gephyrin Phosphorylation and GABAAR Trapping at Inhibitory Synapses. iScience 22, 453–465 (2019).

46. Heine M, et al. Surface mobility of postsynaptic AMPARs tunes synaptic transmission. Science 320, 201–205 (2008).

47. Lois C, Hong EJ, Pease S, Brown EJ, Baltimore D. Germline transmission and tissue-specific expression of transgenes delivered by lentiviral vectors. Science 295, 868–872 (2002).

48. de Chaumont F, et al. Icy: an open bioimage informatics platform for extended reproducible research. Nat Methods 9, 690–696 (2012).

49. De Zitter E, et al. Mechanistic investigation of mEos4b reveals a strategy to reduce track interruptions in sptPALM. Nature Methods 16, 707–710 (2019).

50. Gosse C, Strick TR, Kostrz D. Molecular scaffolds: when DNA becomes the hardware for single-molecule investigations. Curr Opin Chem Biol 53, 192–203 (2019).

51. Keppler A, Gendreizig S, Gronemeyer T, Pick H, Vogel H, Johnsson K. A general method for the covalent labeling of fusion proteins with small molecules in vivo. Nat Biotechnol 21, 86–89 (2003).

52. Gautier A, et al. An engineered protein tag for multiprotein labeling in living cells. Chem Biol 15, 128–136 (2008).

53. Bedet C, et al. Regulation of gephyrin assembly and glycine receptor synaptic stability. J Biol Chem 281, 30046–30056 (2006).

54. Maric HM, et al. Gephyrin-binding peptides visualize postsynaptic sites and modulate neurotransmission. Nat Chem Biol 13, 153–160 (2017).

55. Strick TR, Allemand JF, Bensimon D, Bensimon A, Croquette V. The elasticity of a single supercoiled DNA molecule. Science 271, 1835–1837 (1996).

56. Gosse C, Croquette V. Magnetic tweezers: micromanipulation and force measurement at the molecular level. Biophys J 82, 3314–3329 (2002).

57. Yu Z, et al. A force calibration standard for magnetic tweezers. Rev Sci Instrum 85, 123114 (2014).

58. Ruiz-Gutierrez N, Rieu M, Ouellet J, Allemand JF, Croquette V, Le Hir H. Novel approaches to study helicases using magnetic tweezers. Methods Enzymol 673, 359–403 (2022).

59. Duboc C, Fan J, Graves ET, Strick TR. Preparation of DNA Substrates and Functionalized Glass Surfaces for Correlative Nanomanipulation and Colocalization (NanoCOSM) of Single Molecules. Methods Enzymol 582, 275–296 (2017).

60. Revyakin A, Ebright RH, Strick TR. Single-molecule DNA nanomanipulation: improved resolution through use of shorter DNA fragments. Nat Methods 2, 127–138 (2005).

